# A computational modeling of primary-microRNA expression

**DOI:** 10.1101/2022.04.04.486960

**Authors:** Hansi Zheng, Saidi Wang, Xiaoman Li, Haiyan Hu

**Affiliations:** Department of Computer Science, University of Central Florida, Orlando, Florida, United States of America; Burnett School of Biomedical Science, College of Medicine, University of Central Florida, Orlando, Florida, United States of America

## Abstract

MicroRNAs (miRNAs) play crucial roles in gene regulation. Most studies so far focus on mature miRNAs, which leaves many gaps in our knowledge in primary miRNAs (pri-miRNA). To fill these gaps, we attempted to model the expression of pri-miRNAs in 1829 primary cell types and tissues in this study. We demonstrated that the expression of their associated mRNAs could model the expression of the pri-miRNAs well. These associated mRNAs are different from their corresponding target mRNAs and are enriched with specific functions. The majority of the associated mRNAs of a miRNA are shared across conditions, although a fraction of the associated mRNAs are condition-specific. Our study shed new light on the understanding of miRNA biogenesis and general gene transcriptional regulation.

## Introduction

It is important to study the expression of primary microRNAs (pri-miRNAs). MiRNAs are short endogenous non-coding RNAs. Their biogenesis starts from the transcription of pri-miRNAs, which are processed into precursor miRNAs and eventually become mature miRNAs of ~22 nucleotides (nt) long [1, 2]. The mature miRNAs, usually referred to as miRNAs, bind to their target mRNAs to regulate the target gene expression by degrading the target mRNAs or preventing them from being translated into proteins. The pri-miRNAs are thus the first product of the miRNA biogenesis, which affect the production of the mature miRNAs and the activity of the majority of protein-coding genes the mature miRNAs regulate. Moreover, the expression of pri-miRNA is known to be quite different from that of the corresponding mature miRNAs due to several steps in miRNA biogenesis that process pri-miRNA to mature miRNAs [3]. In fact, previous studies showed that the expression of mature miRNAs did not correlate well with that of precursor miRNAs, and even the mature miRNAs generated from the same primary miRNA transcript had different expression levels [3, 4]. To understand miRNA-involved gene regulation, it is thus indispensable to study the expression of pri-miRNAs.

Most studies so far focus on mature miRNAs [5–12]. The expression of mature miRNAs is routinely measured by small RNA-seq experiments [9]. The co-measurement strategy, which profiles the expression of both mRNAs and miRNAs under related conditions, is commonly employed to pinpoint miRNA target genes [6, 13, 14]. The large-scale analysis of miRNAs and mRNAs gene expression is widely used to define miRNA targets, miRNA modules, miRNA-mRNA co-expression networks, gene regulatory networks, etc. [7–9, 12–18]. Through these studies, it is known that miRNA expression often correlates well with their target mRNAs expression.

Despite many studies on mature miRNAs, the study of pri-miRNA expression is scarce. This scarcity is largely due to the fact that rarely are the pri-miRNAs known, including the miRNA gene transcriptional start sites (TSSs) [19]. Since the TSS of a miRNA gene can be tens or hundreds of thousand base pairs away from the location of the precursor and mature miRNAs, although next-generation sequencing technologies can faithfully identify TSSs of various types of genes, including miRNA genes, it is non-trivial to connect a TSS with a miRNA gene [19–23]. In fact, the majority of the annotated pri-miRNA TSSs were not consistent across studies [19, 22]. To make it even more challenging, pri-miRNAs usually have low and condition-specific expression, short life span, alternative TSSs under different physiological conditions, etc. It is thus no wonder that only a couple of studies have profiled primary miRNA gene expression so far, and it is unclear whether the profiled is truly the pri-miRNA expression [4, 21, 22].

To fill this gap in miRNA studies, we modeled the expression of pri-miRNA in 1829 primary cell types and tissue samples [24]. We found that the expression of the associated mRNAs of pri-miRNAs could reliably model the expression of the pri-miRNAs. These associated mRNAs of a miRNA were in general not the target genes of this miRNA. For a miRNA, its associated mRNAs were enriched with specific functions and mostly conserved across samples, although a small fraction of its associated genes were condition-specific. Our study shed new light on the expression of pri-miRNAs.

## Material and Methods

### 1829 Cap Analysis of Gene Expression (CAGE) samples for expression analysis

We downloaded the normalized expression in 1829 primary cell types and tissues from FANTOM (https://fantom.gsc.riken.jp/5/datafiles/latest/extra/CAGE_peaks/hg19.cage_peak_phase1and2combined_tpm_ann.osc.txt.gz) [24]. We considered CAGE data instead of other TSS-seq data because the CAGE data had the largest number of samples from the same study, which avoided many unexpected discrepancies among samples from different labs. These CAGE data were normalized by the previous study [24]. To determine the expression level of a miRNA or an mRNA, we used the normalized CAGE expression from all peaks located in the corresponding miRNA or mRNA TSS region. When multiple CAGE peaks occurred in a TSS region, we used the expression of the peak with the maximum expression value to represent the expression value of this TSS region.

### The robust set of miRNA TSSs

We previously collected annotated TSSs for 330 miRNAs from fourteen studies [19]. These TSSs were consistently annotated in at least four of fourteen previous studies. Since one miRNA may have multiple annotated TSSs, we considered each annotated TSS as a particular miRNA and actually considered 369 miRNAs. We then calculate the expression of the 369 miRNAs as mentioned above in the 1829 samples. We filtered miRNAs with zero expression in more than 80% of the sample to reduce noise. In this way, 278 miRNAs were remained in our analysis after filtering.

### The consistent set of mRNAs

Since we measured gene expression from the 1829 CAGE experiments, we wanted the expression of mRNAs to be consistent between RNA-seq experiments and the CAGE experiments. In this way, we hoped that the expression of pri-miRNAs we modeled was the expression we normally observed in RNA-seq experiments instead of only in the CAGE experiments.

To define such a consistent set of mRNAs, we used the 22 tissue samples where both RNA-seq and CAGE data were generated (https://www.ebi.ac.uk/arrayexpress/experiments/E-MTAB-1733/samples/?s_page=1&s_pagesize=500) [25]. The CAGE data and the RNA-seq were processed as previously [25] to obtain the gene expression levels for all 27,493 GENCODE annotated mRNA transcripts. We then calculated Spearman’s correlation coefficient for all transcripts across the 22 samples. The transcripts with a correlation larger than 0.75 were chosen as the consistent mRNAs. For a gene with multiple consistent transcripts, we chose the transcript with the largest correlation to represent this gene and filtered all other transcripts. In this way, we obtained 2312 mRNA transcripts. These mRNAs together with their annotated TSSs were used to model the expression of pri-miRNAs.

### The least absolute shrinkage and selection operator (LASSO) regression

We modeled the pri-miRNA expression by LASSO. LASSO is widely used to model gene expression and select variables previously [26–29]. We use the LASSO tool from scikit-learn (https://scikit-learn.org/0.24/, version 0.24.2). The goal of LASSO is to minimize:

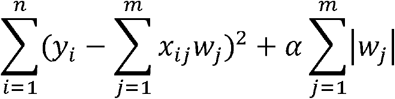

Here the expression of pri-miRNAs in the *i-th* sample was considered as the dependent variables *y*_*i*_, which was a vector of 278 dimensions. The expression of the *j-th* mRNAs in the *i-th* sample was considered as the independent variables *x*_*ij*_. The *w*_*j*_ was the coefficient vector of 278 dimensions to describe the importance of the *j-th* mRNA to the pri-miRNAs. The LASSO method tried to optimize the above function by making certain *w*_*j*_ to be zero and choosing the remaining mRNAs as the associated mRNAs. Both the values of the dependent and independent variables were normalized to have a mean zero and standard deviation one before the LASSO regression was applied.

We considered three different neighborhood sizes, 100, 300, and 500 base pairs, around the TSSs to measure the expression of pri-miRNAs and mRNAs. For a given neighborhood size, the expression of a pri-miRNA and a mRNA was calculated as the normalized expression of the CAGE peak located in the corresponding neighborhood. If multiple CAGE peaks were in the neighborhood of a TSS, we used the largest expression value of these peaks. We then applied the LASSO regression to the data for a given neighborhood size. Because the neighborhood size did not affect the model much, we presented the results from the neighborhood size of 100 base pairs.

To measure how well the expression of pri-miRNAs was modeled, we calculated the correlation coefficient of the predicted expression of a pri-miRNA with its actual expression. We calculated both Pearson’s correlation and Spearman’s correlation per miRNA and per sample. For the per miRNA correlation, we considered the two vectors of expression values across 1829 samples for a miRNA. For the per sample correlation, we considered the two vectors of expression values across 278 miRNAs for a sample. The significance of a correlation *r* was approximated by the t-test p-values 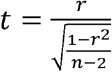, which asymptotically followed a t-distribution with the degree *n-2*, with *n=278* for the per-sample correlations and *n=1829* for the per-gene correlations.

### The GO analysis

We inferred the significantly enriched GO terms for the associated mRNAs of a miRNA. The gene symbols of the 2312 consistent mRNAs were considered as the population of all genes. The associated mRNAs for each miRNA were served as the target list of genes. We then searched for the enriched GO terms in every target list of genes compared with the population of all genes using the GOrilla tool [30]. We recorded the number of GO terms identified by GOrilla for each miRNA with the false discovery rate cutoff 1.

## Results

### The expression of the pri-miRNAs was reliably predicted

We modeled the expression of pri-miRNA in 1829 primary cell types and tissues (Material and Methods). In brief, we considered the TSSs of a miRNA consistently annotated in at least four of fourteen previous studies as the TSS of this miRNA [19]. We then measured the expression of this miRNA by the relative log expression normalized counts of reads mapped to the neighborhood of the TSS in the Cap Analysis of Gene Expression (CAGE) experiments [24]. Finally, we modeled the expression of the pri-miRNAs with the expression of mRNAs in these 1829 CAGE experiments by the least absolute shrinkage and selection operator (LASSO) regression [31]. LASSO selected a subset of mRNAs for each miRNA. We called these selected mRNAs for a miRNA as its associated mRNAs.

We found that the expression of the associated mRNAs could reliably model the expression of pri-miRNAs (Figure 1A). We calculated the correlation of the predicted expression value by the LASSO model with the true expression value. The minimum, mean, and median Pearson’s correlation per miRNA was 0.59, 0.91 and 0.91, respectively (p-value = 0 for all correlations). Similar, the minimum, mean and median Spearman’s correlation per miRNA was 0.27, 0.77 and 0.80, respectively (p-value = 0 for all correlations). If we measured the similarity of the predicted expression in every sample, the minimum, mean, and median Pearson’s correlation per sample was 0.51, 0.87 and 0.89, respectively (p-value = 0 for all correlations). Correspondingly, the minimum, mean and median Spearman’s correlation per sample was 0.43, 0.81 and 0.82, respectively (p-value < 3.12e-14 for all correlations). The significant correlation suggested that the expression of the associated mRNAs could reliably model the pri-miRNA expression.

**Figure 1.**
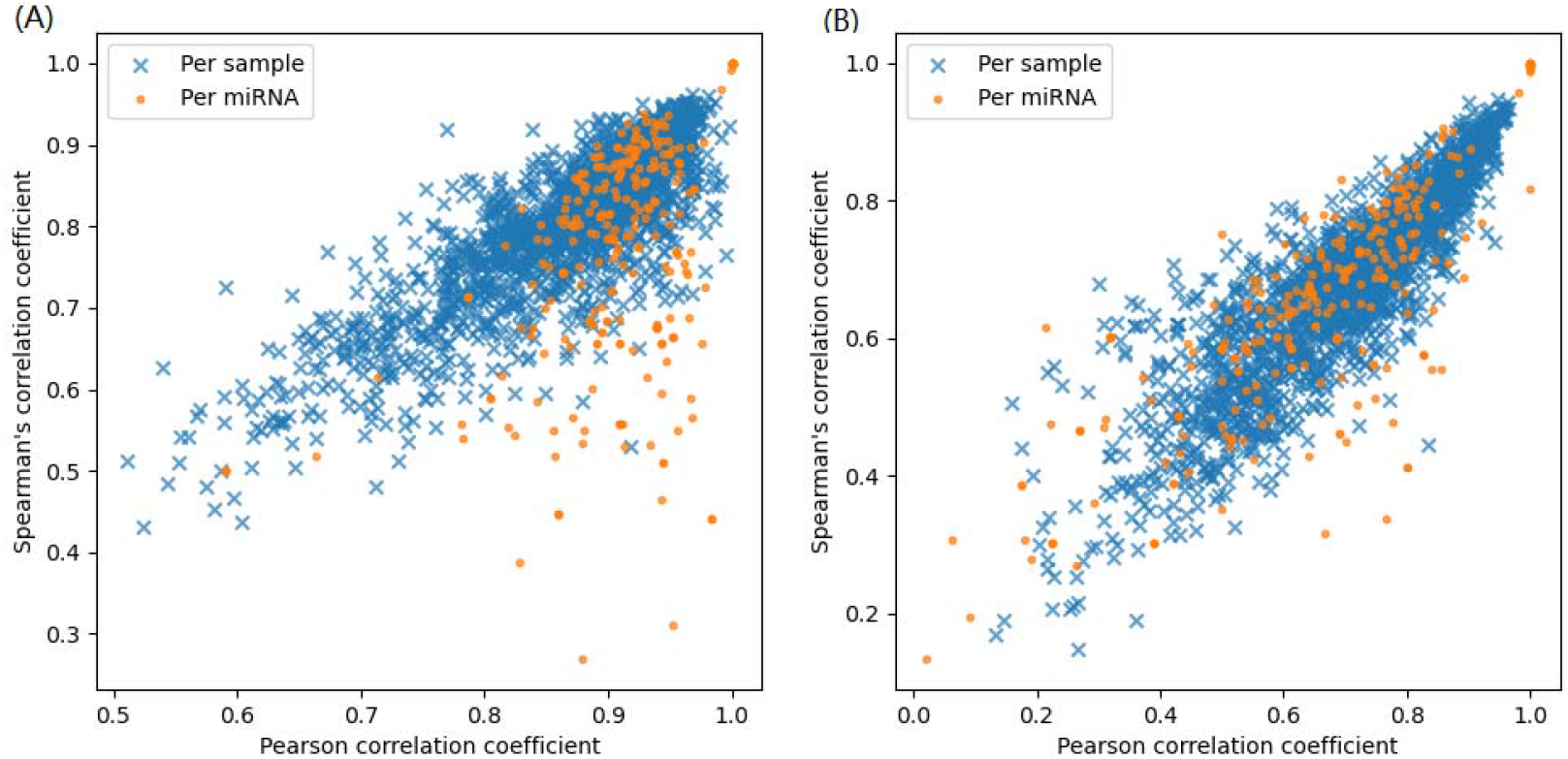
(A)The correlation coefficient between the predicted expression and the true expression in all samples; (B) correlation coefficient of 10-fold cross validation between the predicted expression and the true expression value.

We further examined the miRNAs with their expression accurately predicted (Pearson’s correlation >0.9) and the miRNAs with the expression not predicted so well (Pearson’s correlation <0.6) (Table 1). Note that the correlation was larger than 0.59 for all miRNAs for the two types of correlation calculations. We found that the miRNAs with their expression accurately predicted were pri-miRNAs with much higher expression levels and much larger expression variation. On the contrary, miRNAs that were not modeled so well were pri-miRNAs with low expression and low expression variation. For instance, the miRNAs modeled well had a maximum expression value and standard deviation of 313.46 and 427.57, while the miRNAs modeled not so well had the corresponding value as 0.20 and 0.43, respectively. In fact, for every miRNA modeled not so well, they had zero expression in at least 65.12% of the samples. In other words, these miRNAs were not modeled so well because they were not so related to the experimental conditions these samples considered. If we excluded the miRNAs with their expression standard deviation smaller than 3 in these 1829 samples, the Pearson’s correlation of the predicted expression with the true expression was larger than 0.83 for each of the remaining 240 miRNAs (p-value = 0). We called these 240 miRNAs active miRNAs in the 1829 samples.

**Table 1.** The minimum, maximum, and median of Pearson’s correlation coefficient of three groups of miRNAs. The miRNAs in the high group had a correlation > 0.9. The miRNAs in the low group had a correlation < 0.6. The miRNAs in the active group had the standard deviation of expression larger than three.

| **Group** | **Min**<br>**correlation** | **Max**<br>**correlation** | **Median**<br>**correlation** | **Min**<br>**expression** | **Max**<br>**expression** | **Median**<br>**expression** |
| --- | --- | --- | --- | --- | --- | --- |
| **High** | 0.90 | 1.00 | 0.94 | 0.37 | 313.48 | 16.93 |
| **Low** | 0.59 | 0.59 | 0.59 | 0.13 | 0.13 | 0.13 |
| **Active** | 0.83 | 1.00 | 0.92 | 0.47 | 313.48 | 15.70 |

The good modeling of the expression of the 240 pri-miRNAs suggested that the expression of almost all pri-miRNAs could be predicted well by their associated mRNAs when related samples were considered.

The above analysis was based on all 1829 samples. To see whether the LASSO models trained on a subset of samples could predict pri-miRNA expression on the remaining samples, we did ten-fold cross-validation (Figure 1B). On average, the minimum, mean and median Pearson’s correlation per miRNA was 0.02, 0.65 and 0.67, respectively. The minimum, mean and median Spearman’s correlation per miRNA was 0.13, 0.65 and 0.65, respectively. Correspondingly, the minimum, mean and median Pearson’s correlation per sample was 0.13, 0.70, and 0.67, respectively. The minimum, mean and median Spearman’s correlation per sample was 0.15, 0.68 and 0.69, respectively. Although lower compared with the model trained with all 1829 samples, the correlation was still significantly large (p-value of the mean and median correlation was 0), suggesting that the pri-miRNA expression was reliably predicted. The lower correlation in the cross-validation also indicated that the regulation of the pri-miRNA expression was sample-specific, and the inferred model based on a specific subset of samples would predict pri-miRNA expression better for that particular subset of samples.

### The associated mRNAs were different from target mRNAs

Previous studies modeled mature miRNA expression with the expression of their target mRNAs [8, 13]. The target mRNAs of a miRNA contain its target sites and can be bound by this miRNA. We thus compared the associated mRNAs inferred above with the target mRNAs for each miRNA. The target mRNAs of a miRNA were retrieved from two sources (Material and Methods). One was the computationally predicted targets by the TargetScan Version 7.2 tool [5]. The other was the annotated miRNA targets from the miRTarBase Version 8.0 database [32].

We found that the associated miRNAs were different from the target mRNAs for every miRNA. On average, only about 2.22% of the associated mRNAs were the targetScan mRNAs, and about 5.26% of targetScan mRNAs were the associated mRNAs for a miRNA. Similarly, the corresponding percentage was 1.06% and 5.56%, respectively, when the miRTarBase mRNAs were compared with the associated mRNAs for every miRNA. The rare overlap between the two types of mRNAs may make sense, since if the associated mRNAs were largely the target mRNAs, the pri-miRNAs would have a highly correlated expression with their mature miRNAs, which was not the case in previous studies [3, 4]. The rare overlap thus corroborates the previous observations that pri-miRNAs and mature miRNAs have in general uncorrelated expression patterns [3, 4].

Since the associated mRNAs modeled the pri-miRNA expression better than the target mRNAs, we hypothesized that their expression correlated better with the expression of the corresponding pri-miRNA than the expression of the target mRNAs. We found that for a miRNA, the expression of most of its associated mRNAs indeed correlated better with its expression than the expression of its target mRNAs across the 1829 samples, no matter whether the target mRNAs were defined by targetScan or miRTarBase (Figure 2). When we compared the expression correlation of the associated mRNAs with the expression correlation of the target mRNAs, we found that 84.53% of miRNAs had a significantly higher correlation with their associated mRNAs than with their target mRNAs (Mann-Whitney p-value < 0.001).

**Figure 2.**
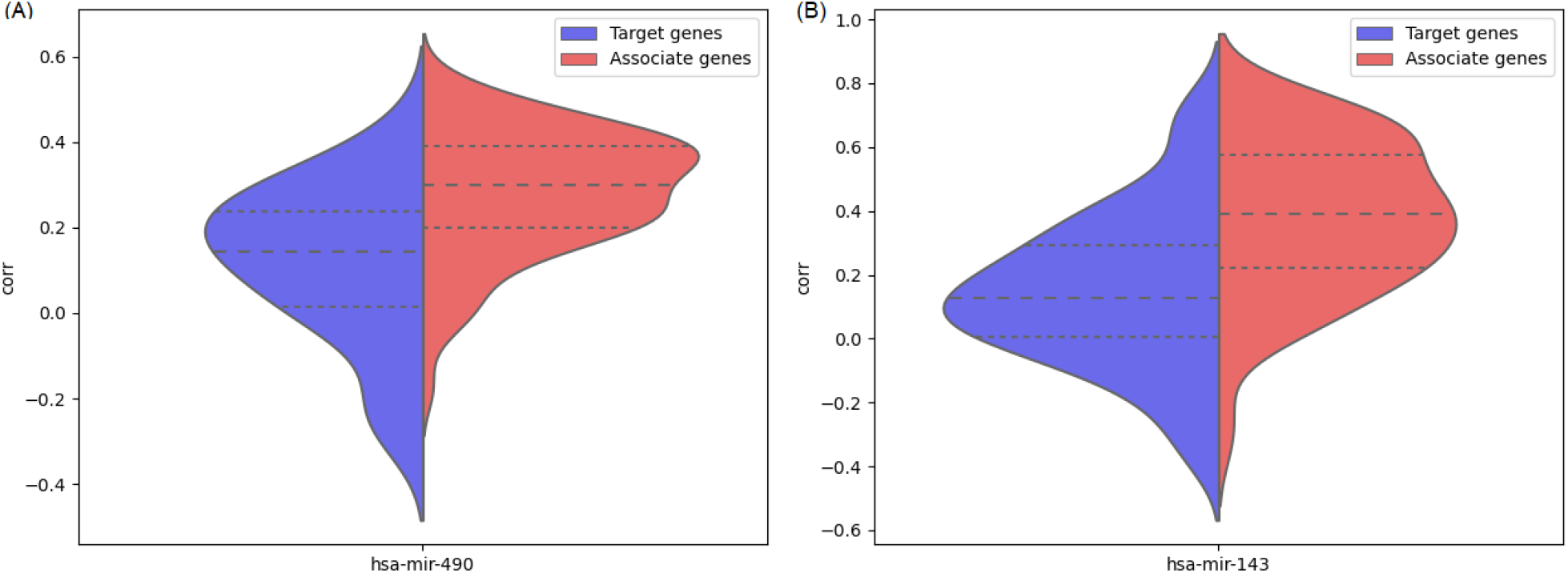
Spearman’s correlation of target genes from miTarBase and TargetScan compared with the Spearman’s correlation of associate genes (A). has-miR-490; (B) hsa-mir-143.

Despite the rare overlap of the two types of mRNAs for most miRNAs, two miRNAs, has-mir-326 and hs-mir-454, both had seven associated mRNAs overlapped with the targetScan mRNAs. For these two miRNAs, their associated mRNAs significantly overlapped with their targetScan mRNAs (hypergeometric testing p-value 0.026 and 0.006). Interestingly, the miRNA mir-454 also had five associated mRNAs overlapped with the miRTarBase mRNAs (hypergeometric testing p-value 0.001).

### The associated mRNAs were biologically sound

The above analysis showed that the associated mRNAs had more correlated expression pattern with the pri-miRNAs than the target mRNAs, implying the functionality of these associated mRNAs for each miRNA. We further investigated other properties of the associated mRNAs. We found that most of them were not affected by the neighbor sizes around miRNA TSSs and different samples used. Moreover, these associated mRNAs for most miRNAs had enriched functional annotations.

We checked how different neighborhood sizes might affect the associated mRNAs inferred for a miRNA (Material and Methods). We considered 100, 300 and 500 base pairs around an annotated miRNA TSS as the TSS regions to measure the normalized gene expression of miRNAs and mRNAs. For the three neighborhood sizes, the median number of associated mRNAs identified was 164, 163, and 164, respectively, indicating the robustness of the inferred associated mRNAs. We also found that more than 77.0% of the associated mRNAs were the same for a miRNA when different neighborhood sizes were used, suggesting that the associated mRNAs were likely to be biologically meaningful and intrinsically related to the corresponding miRNAs. Although most associated mRNAs were the same for a miRNA, a fraction of them were different, most likely due to the condition-specific expression of the pri-miRNAs.

We also studied how different samples may change the associated mRNAs inferred for a miRNA. For a given miRNA, we compared its associated mRNAs inferred from each of the ten folds in the ten-fold cross-validation with the associated mRNAs inferred with all samples (Figure 3A). Here the neighborhood size was set to be 100 base pairs since different neighborhood sizes did not change the predictions much. Interestingly, the median number of the associated mRNAs was still around 163. Moreover, on average, about 80.79% of the associated mRNAs were shared between every fold and the model with all samples. We also found 79.78% of the associated genes were shared between every fold and at least four other folds (Figure 3B). Together with the above analysis, it was evident that most associated mRNAs were the same when different neighborhood sizes or different subsets of samples were used. It also showed that a fraction of the associated mRNAs were condition-specific, occurring in specific sets of samples.

**Figure 3.**
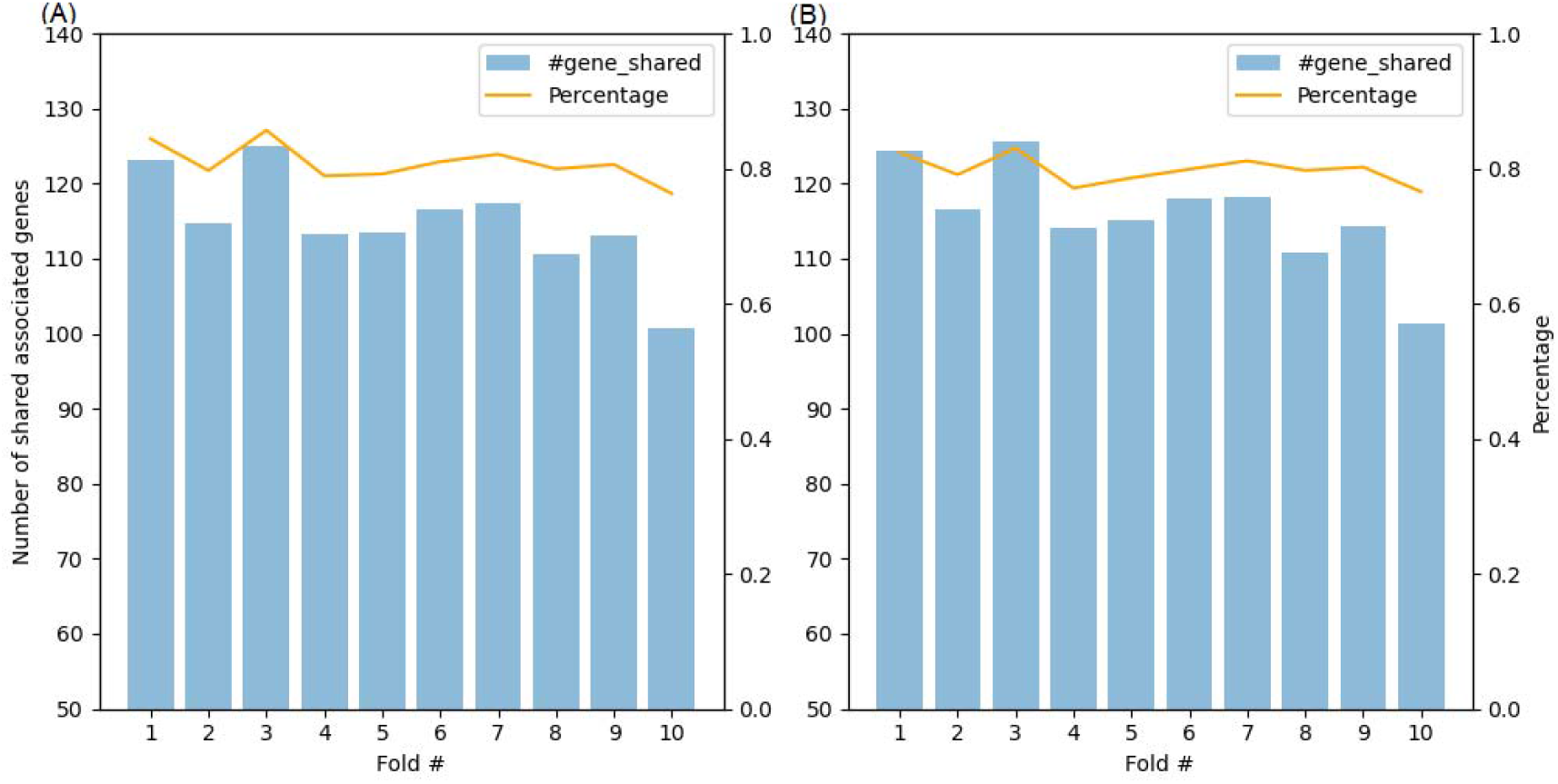
(A) The number and percentage of associated mRNAs in each fold shared by the full model with all samples. (B) The number and percentage of associated mRNAs in each fold shared by at least four other folds.

With the associated mRNAs for each miRNA, we investigated whether they significantly shared gene ontology (GO) functions [33]. With the false discovery rate cutoff 1, more than 58.63% pri-miRNAs had at least one GO term significantly shared by its associated mRNAs. Among these pri-miRNAs, the median number of significantly shared GO terms was three. The GO analysis suggested that the associated mRNAs were likely biologically meaningful. Note that not all miRNAs had their associated mRNAs significantly shared GO functions, partially due to the imperfect GO annotation.

## Discussion

The study of pri-miRNA expression is still in its infancy. Here we modeled the pri-miRNA expression in 1829 primary cells and tissues. We demonstrated for the first time that the expression of the associated mRNAs could reliably predict the expression of the pri-miRNAs. These associated mRNAs barely overlapped with their target mRNAs while having a more correlated expression with the pri-miRNAs than the target mRNAs. For most miRNAs, their associated mRNAs significantly shared GO functions.

We demonstrated that the expression of almost all pri-miRNAs could be reliably modeled. In fact, the predicted expression had a correlation of more than 0.82 with the actual expression for all pri-miRNAs with the standard deviation of the expression larger than three in the 1829 samples (p-value=0, Table 1). We showed that the miRNAs modeled not so well were likely to have low expression and low expression variation. With more samples available in the future, one could model the expression of more pri-miRNAs.

More than a dozen studies previously predicted or annotated miRNA TSSs [19, 22, 34–37]. These predicted or annotated TSSs were often inconsistent between different studies [19, 22]. A recent survey identified 369 miRNA TSSs consistent in at least four of the fourteen previous studies for 330 miRNAs [19]. We selected 278 of these 369 miRNA TSSs that showed transcriptional activities in at least 20% of the 1829 samples in this study. Although we did not consider the alternative miRNA TSSs, these miRNA TSSs were likely the best set we could have currently, since they were shown to have better qualities previously [19, 22].

In addition to the 278 miRNA TSSs, we considered 2312 mRNAs to model the expression of pri-miRNAs. Because we measured gene expression through the CAGE data, we narrowed our analysis to the 2312 mRNAs that had consistent expression patterns in CAGE and RNA-seq experiments. Moreover, we applied LASSO to model the expression of pri-miRNAs, which was thus more likely to capture only linear relationships between the miRNAs and the mRNAs. In the future, with more accurate annotation of miRNA TSSs and a better understanding of TSS-seq data, more sophisticated approaches and more comprehensive studies can be carried out to involve more pri-miRNAs and more mRNAs [38–40].

